# Identification of a new gene *Rf9* and unravelling the genetic complexity for controlling fertility restoration in hybrid wheat

**DOI:** 10.1101/2020.06.20.162644

**Authors:** Fahimeh Shahinnia, Manuel Geyer, Annette Block, Volker Mohler, Lorenz Hartl

## Abstract

Wheat (*Triticum aestivum L.*) is a self-pollinating crop whose hybrids offer the potential to provide a major boost in yield. Male sterility induced by the cytoplasm of *Triticum timopheevii* is a powerful method for hybrid seed production. Hybrids produced by this method are often partially sterile and full fertility restoration is crucial for wheat production using hybrid cultivars. To identify genetic loci controlling fertility restoration in wheat, we produced two CMS-based backcross (BC_1_) mapping populations. The restorer lines Gerek 79 and 71R1203 were used to pollinate the male-sterile winter wheat line CMS-Sperber. Seed set and numbers of sterile spikelets per spike were evaluated in 340 and 206 individuals of the populations derived from Gerek 79 and 71R1203, respectively. Genetic maps were constructed using 930 and 994 SNPs, spanning 2,160 and 2,328 cM over 21 linkage groups in the two populations, respectively. Twelve quantitative trait loci (QTL) controlled fertility restoration in both BC_1_ populations, including a novel *restorer-of-fertility (Rf)* locus flanked by the single nucleotide polymorphism (SNP) markers *IWB72413* and *IWB1550* on chromosome 6AS. The locus was mapped as a qualitative trait in the BC_1_ Gerek 79 population and was designated *Rf9.* Ninety-three putative candidate genes were predicted for the QTL region on chromosome 6AS. Among them were genes encoding tetratricopeptide and pentatricopeptide repeat-containing proteins in rice known to be associated with fertility restoration. This finding is a promising step to better understand the functions of genes for improving hybrid wheat.

## Introduction

Since the discovery of male sterility and restoration systems in the 1960s, hybrid wheat triggered attention due to its potential for improved grain and straw productivity and yield stability particularly under harsh and marginal environments (Longin et al. 2012; Whitford et al. 2013). The major gains of hybrid versus line varieties are improved trait values due to heterosis (Castillo et al. 2014). Heterosis or hybrid vigor is the phenomenon by which the progeny derived from a cross of two inbred lines outperform the parent lines and has provided considerable economic benefits in worldwide crop production (Chen and Liu 2014). Heterosis in winter wheat has been reported to provide uniform plant establishment and tolerance against frost, lodging, and diseases such as leaf rust, stripe rust, *Septoria tritici* blotch and powdery mildew (Gupta et al. 2019).

To harness yield gains associated with hybrid vigor, the cytoplasmic male sterility (CMS) system provides a cost-effective tool for efficient hybrid seed production (Chen and Liu 2014). CMS in plants is based on rearrangements of mitochondrial DNA that lead to chimaeric genes and a condition under which a plant is unable to produce fertile pollen (Whitford et al. 2013). CMS evades the need for manual removal of anthers, thus facilitating a technology to produce unlimited numbers for hybrid plants. It has been successfully used in crops such as rye, rice, maize and sunflower (Castillo et al. 2015). In wheat, the outcome of a hybridization event involving *T. timopheevii* as the female parent and *Triticum aestivum* (Bohra et al. 2016) causes CMS. The use of *T. timopheevii* cytoplasm in bread and durum wheat creates male sterility, whereas female fertility is not impaired (Lukaszewski 2017). CMS-based hybrid production uses three different breeding lines, a CMS line, a maintainer line, and a restorer line (Chen and Liu 2014). The CMS line is used as the female parent with at least one CMS-causing gene in the *T. timopheevii-derived* cytoplasm and lacking functional nuclear-encoded *Restorer-of-fertility (Rf)* genes (Schnable and Wise 1998). The maintainer line serves as the male parent in crosses for the propagation and maintenance of the CMS line, with the same nuclear genome as the CMS line, but a normal fertile *T. aestivum* cytoplasm. The restorer line retains (a) functional *Rf* gene(s) and acts as the male parent to cross with the CMS line to produce the F1 hybrid seeds. In F1 plants, the *Rf* gene(s) restore(s) male fertility, and the combination of the nuclear genomes from the CMS line and the restorer line produces hybrid vigor (Chen and Liu 2014). For commercial hybrid seed production, a male-sterile line has to be crossed with a line carrying dominant restorer alleles and suitable pollinator qualities (Whitford et al. 2013). While fertility restoration is a crucial trait in hybrid breeding, hybrids produced using this method are often partially sterile due to the complex interaction between mitochondrial and nuclear genes controlling male specificity, occurrence, and restoration of fertility (Chen and Liu 2014). Therefore, incomplete fertility restoration poses a major bottleneck for hybrid wheat breeding, as it impairs the effect of heterosis on grain yield and the uniformity or quality of end-use products (Keydel 1973; Geyer et al. 2018).

Fertility restoration is a genetically complex process and is mainly controlled by the mitochondrial genome in interaction with *Rf* genes (Castillo et al. 2014). Besides, it is known that fertility restoration is influenced by environmental factors including photoperiod, water stress, light intensity and temperature (Castillo et al. 2015). The mitochondrial gene families produce proteins that share the common structural organization of similar repeated helical motifs and include half-tetratricopeptides, pentatricopeptide repeat (PPR) proteins, and mitochondrial transcription termination factors (mTERF) (Pan et al. 2019). Tetratricopeptide repeat (TPR) motifs occur in proteins from bacteria to humans and play important roles in various processes, including hormone regulation, protein kinase inhibition, protein folding, cell cycle and regulation of reproductive development (Yu et al. 2016). The effect of cytoplasmic male sterility can be weakened by inhibiting the accumulation of the CMS-conferring gene products through the function of a class of *Rf* genes, generally belonging to a large family of genes that encode organelle-targeted PPR proteins (Hu et al. 2012). mTERF genes are widely distributed in metazoans, plants and green alga. They regulate transcription, translation and DNA replication of mitochondrial genes in metazoans while regulating gene expression in chloroplasts and mitochondria in plants (Pan et al. 2019). In wheat, the presence of eight major genes *(Rf1-Rf8)* for timopheevii-based cytoplasmic male sterility is known and assigned to the chromosomes 1A, 7D, 1B, 6B, 6D, 5D, 7B and 2D, respectively (Mukai and Tsunewaki 1979; Sinha et al. 2013; Gupta et al. 2019). *Rf1* and *Rf3* are the most effective genes for achieving restoration in wheat (Geyer et al. 2016, 2018; Würschum et al. 2017). Previous studies have indicated that combinations of two or three major *Rf*genes and restorer genes with small effect or low penetrance (modifier loci) can modify the degree of fertility restoration (Ma et al. 1995; Ahmed et al. 2001; Zhou et al. 2005; Stojałowski et al. 2013; Whitford et al. 2013; Lukaszewski 2017). Consequently, attempts are made to pyramid multiple dominant or partially dominant alleles of the most favourable genes or quantitative trait loci (QTL) including those involved in epistatic interactions in order to achieve complete fertility restoration in hybrid wheat (Zhao et al. 2013; Jiang et al. 2017; Gupta et al. 2019). Understanding the genetic mechanisms underlying restoration of fertility and developing elite restorer lines are crucial to overcoming the intricate barriers in hybrid breeding programs. For this reason, our objective was to identify new genetic loci controlling fertility restoration that can be employed in hybrid wheat breeding. Here, we developed two CMS-based backcross (BC) mapping populations, which we used for QTL mapping and identification of candidate genes. Our study identified a new *Rf* gene *(Rf9)* and novel QTL for seed set on chromosomes 4AL, 5BL and 6AS and for number of sterile spikelets per spike on chromosomes 1DS, 2AL and 6AS. Putative candidate genes located in the target regions are discussed.

## Materials and methods

### Plant materials and population development

Two BC_1_ mapping populations were developed using Gerek 79 and 71R1203 as fertility-restoring parental lines. The winter wheat cultivar Gerek 79 (PI 559560, pedigree: Mentana/Mayo-48//4-11/3/Yayla-305) originated in 1979 from the Transitional Zone Agricultural Research Institute, Anadolu ARI, Eskisehir, Turkey (http://wheatpedigree.net/sort/show/21747). The restoration capacity of Gerek 79 was found in initial screening experiments (unpublished) when pollinating CMS-Sperber with cultivars from various regions and testing the hybrids for self-fertility in the greenhouse at the Bavarian State Research Centre for Agriculture (LfL). The winter wheat restorer line 71R1203 (PI 473552, pedigree: NB542437/CI 13438//2*Burt/3/NB542437/2*CI13438) was developed in 1982 by USDA-ARS and Washington State University. The specific fertility restoration genes possessed by each of the two sources had not been previously determined, but 71R1203 was known to potentially carry restorer genes *Rf1* and *Rf2* that are present in NB542437 (Allan and Rubenthaler 1984). The variety Sperber (registered 1983) and CMS-Sperber are maintained at the LfL. Seeds of Gerek 79 were kindly provided by Prof. Friedrich Zeller (Technical University of Munich, Freising, Germany) and are available at the Germplasm Resources Information Network (GRIN), U.S. National Plant Germplasm System. Seeds of line 71R1203 were obtained from National Small Grains Collection, US. Gerek 79 and 71R1203 were used as restorer lines in crosses with the male-sterile winter wheat line CMS-Sperber. The hybrids were then backcrossed with the maintainer line Sperber to develop the mapping populations CMS-Sperber/Gerek 79//Sperber (BC_1_ Gerek 79) and CMS-Sperber/71R1203//Sperber (BC_1_ 71R1203).

### Field trials and phenotyping

BC_1_ Gerek 79 population was vernalised in a climate chamber at 6 °C for 8 weeks and planted in spring 2019 in an LfL field at Freising (48°24’12.64”N, 11°44’55.54”E), Germany. The BC_1_ 71R1203 population was sown in autumn 2018 in the field at KWS LOCHOW GMBH in Bergen (52°48’30.13”N, 9°57’49.46”E), Germany. To assess fertility restoration in the mapping populations, four emerging spikes from the main tillers of each plant were covered before anthesis using glassine bags. After ripening, the spikes were harvested and the seed set (as the restored fertility trait) and number of sterile spikelets per spike (as the non-restored fertility trait) were counted. The seed set of a plant was calculated as the number of kernels divided by the number of spikelets, averaged over all four bagged spikes per individual. Plants were considered fertile if they had at least one seed per spike and male sterile when no seed was produced. Observed ratios of fertile to sterile plants in each mapping population were tested against the expected segregation pattern using *chi-squared goodness*-of-*fit tests*. Statistical analyses including descriptive statistics, correlation and frequency distribution of the traits were conducted in SigmaPlot (Systat Software, San Jose, CA, USA).

### Genotyping and linkage analysis

Genomic DNA of parental lines and BC_1_ progenies was extracted from young leaf tissues following the procedure of Plaschke et al. (1995). Based on fertility restoration phenotypic data, the DNA of 273 and 184 individuals from BC_1_ Gerek 79 and BC_1_ 71R1203, respectively, were selected for genotyping using a bead chip comprising 16,762 Single Nucleotide Polymorphism (SNP) markers selected from the 90K iSelect^®^ array (Wang et al. 2014). SNP genotyping was done by KWS SAAT S & Co, KGaA, Einbeck, Germany. The raw SNP data were analysed as described by Geyer et al. (2018). Briefly, all monomorphic SNPs and those with more than 10 % missing values and a minor allele frequency of less than 10 % were discarded from further analysis using the synbreed package V0.12-6 (Wimmer et al. 2012) in R (R Core Team, 2017). Linkage analysis was done using JoinMap^®^ (Kyazma BV, Wageningen, The Netherlands). The Kosambi mapping function (Kosambi 1944) was used to convert recombination frequencies into centimorgans (cM).

To determine whether Gerek 79 and 71R1203 carried *Rf3,* they were genotyped with SNP marker *IWB72107*, earlier shown to have a high potential for predicting *Rf3* (Geyer et al. 2016). The SNP-containing sequence for *IWB72107* was retrieved from The Triticeae Toolbox (T3, https://triticeaetoolbox.org/wheat/) and converted to a Kompetitive Allele Specific Polymerase (KASP) chain reaction marker assay (Table 1). Plants were genotyped according to the manufacturer’s instructions (LGC Genomics, Hoddeson, UK). Each KASP reaction was prepared in a volume of 10 μL with 5 μL DNA and 5 μL of the genotyping master mix. Amplification was carried out using the CFX96 Touch™ Real-Time PCR SNP Detection System (Bio-Rad, CA, USA), starting with 15 min at 94 °C, followed by 40 cycles of PCR with 94 °C for 20 s and 65 °C for 1 min and 10 cycles of touch down PCR where the annealing temperature was gradually reduced by 0.8 °C per cycle. Endpoint analysis and allelic discrimination related to SNP calls were accomplished using the CFX96 Touch™software (BioRad, CA, USA). The DNA of the restorer line Primepi was used as a reference control for *Rf3* (Geyer et al. 2016).

**Table 1.**
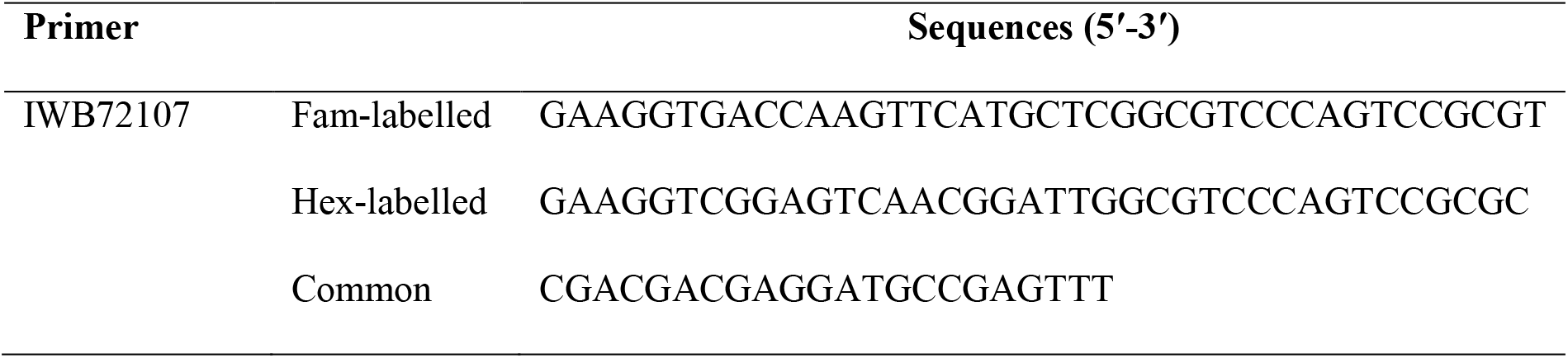
Primer sequences designed for *KASP_IWB72107* marker associated with the gene *Rf3*.

### QTL mapping

To detect QTL controlling seed set and number of sterile spikelets per spike in BC_1_ populations, composite interval mapping with a 5-cM window and a maximum of 10 marker cofactors per model was carried out using Windows QTL Cartographer version 2.5 (Wang et al, 2012). Tests were performed at 1-cM intervals, and cofactors were selected by forward-backward stepwise regression Model 6 (Wang et al. 2012; Shahinnia et al. 2009). Genome-wide, trait-specific threshold values (α = 0.05) of the likelihood ratio test statistic for declaring the presence of a significant QTL was predicted by random sampling of phenotypic data and conducting 2000 permutations test (Churchill and Doerge 1994). The additive effect of an allelic substitution at each QTL and the phenotypic variation explained by a QTL (*R*^2^) conditioned by the composite interval mapping cofactors involved in the model was calculated at the most likely QTL position. The LOD peak of each significant QTL was reflected as the QTL location on the linkage map. To identify markers associated with trait variation located in the confidence interval of a target QTL, single marker analysis was performed using Wald statistics (Kenward et al. 1997; Shahinnia et al. 2016). QTL designation followed the recommended rules for wheat (https://wheat.pw.usda.gov/ggpages/wgc/98/Intro.htm). QTL nomenclature *(Qphenotype.lab-chromosome.Qnumber)* included ‘lfl’, ‘Rf’ and ‘StS’ representing ‘Bayerische Landesanstalt für Landwirtschaft’ (LfL), seed set and number of sterile spikelets per spike, respectively. The *Qnumber* after the chromosome designation refers to overlapping QTL identified on the same chromosome in the two BC_1_ populations.

### Physical mapping and identification of candidate genes

Candidate genes were identified to better understand the genetic basis of fertility restoration. Sequences and physical positions of SNP markers within the QTL regions were retrieved based on the entire bread wheat NRGene genome assembly version 1.0 (https://wheat-urgi.versailles.inra.fr/Seq-Repository/Assemblies) (Appels et al. 2018) using the Triticeae Toolbox (T3, https://triticeaetoolbox.org/wheat/). Candidate genes were identified using web application POPSEQ Ordered *Triticum aestivum* Gene Expression (POTAGE) (Suchecki et al. 2017). The obtained sequences for SNP markers were then used as a query for BLASTn search in POTAGE (http://crobiad.agwine.adelaide.edu.au/potage/). The search by sequence option allowed convenient access to high confidence predictions of genes within the CSS contigs and functional annotation with the respective Munich Information Centre for Protein Sequences (MIPS) (ftp://ftpmips.helmholtz-muenchen.de/plants/wheat/IWGSC/genePrediction_v2.1/), supplemented with rice annotations translated sequence similarities (RGAP version 7). *In silico* expression values for tissue series of the wheat spike, root, leaf, grain and stem organs at different developmental stages (Zadoks et al. 1974) was obtained through the WheatExp (https://wheat.pw.usda.gov/WheatExp/) (Pearce et al. 2015) and a bread wheat tissue series RNA-Seq data set (http://urgi.versailles.inra.fr/files/RNASeqWheat/) in POTAGE. Fragments per kb per million reads (FPKM) were used to show the gene expression quantity, thus avoiding the influence of sequencing length and differences on expression values.

## Results

### Evaluation of fertility restoration

The values for seed set and number of sterile spikelets per spike in BC_1_ Gerek 79 and BC_1_ 71R1203 are presented in Table 2. Whereas a 1:1 segregation ratio for fertile to sterile lines was observed in BC_1_ Gerek 79 (174:166) and BC_1_ 71R1203 (100:106), the average seed set was higher in BC_1_Gerek 79 (0.5) than in BC_1_ 71R1203 (0.3). Number of sterile spikelets per spike showed a negative correlation with seed set in both BC_1_ Gerek 79 (*r* = −0.65) and BC_1_ 71R1203 (*r* = −0.87). Frequency distribution of the traits ranged between 0-2.5 and 0-2.1 for seed set (Fig. 1a, b), whereas a range between 5.5-29.5 and 3.7-23.0 was observed for number of sterile spikelets per spike (Fig. 1c, d) in BC_1_ Gerek 79 and BC_1_ 71R1203, respectively.

**Fig. 1.**
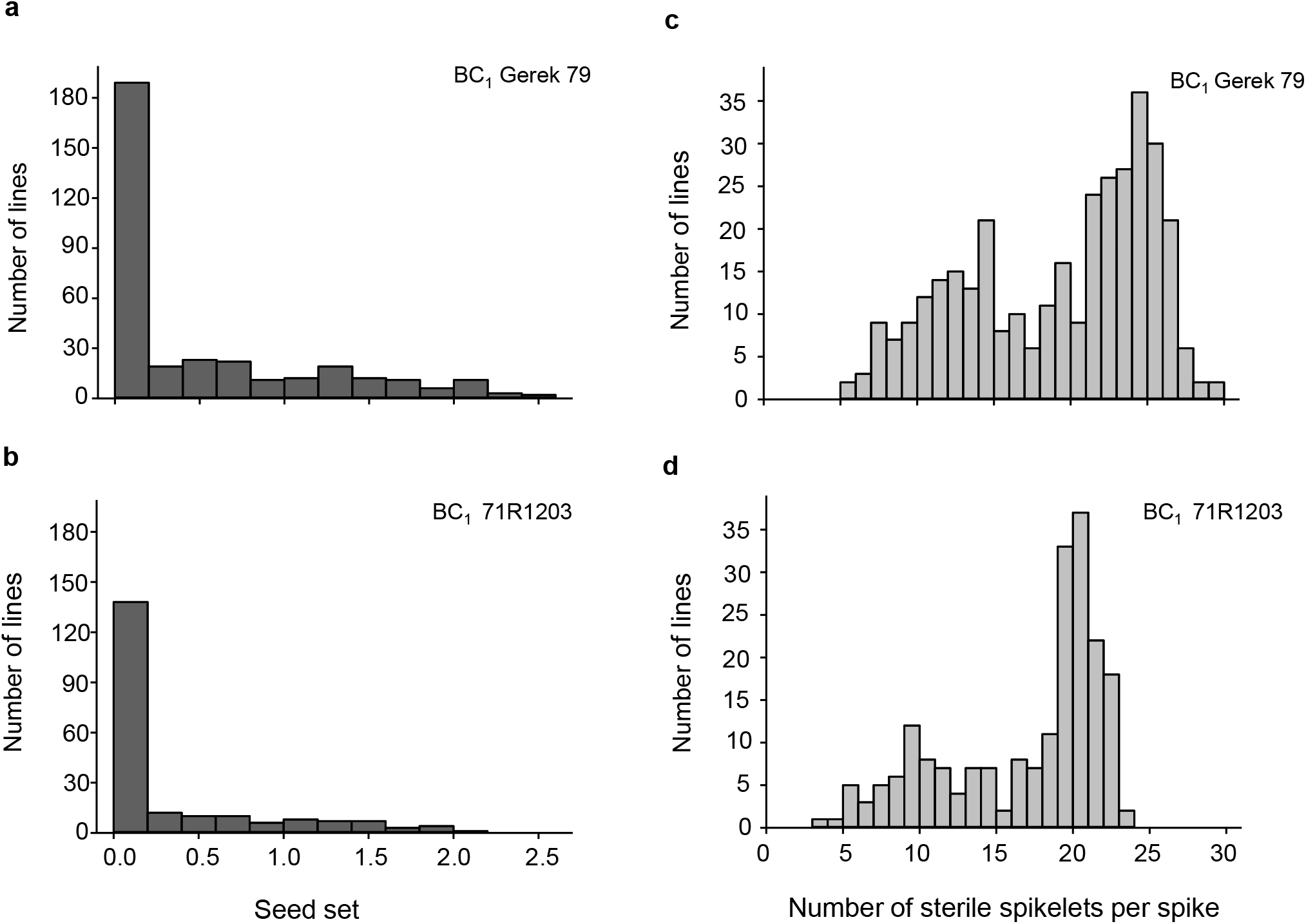
Frequency distribution of phenotypes for (a, b) seed set and (c, d) number of sterile spikelets per spike in BC_1_ Gerek 79 and BC_1_ 71R1203, respectively.

**Table 2.** Descriptive statistics for seed set and number of sterile spikelets per spike in BC_1_ Gerek 79 and BC_1_ 71R1203. Segregation of the number of fertile *vs.* sterile plants was compared to a 1:1 ratio using the *χ*^2^ tests.

| Trait/ Population | No.<br>lines | Mean | Max | Min | SE | No.<br>fertile:sterile | $\chi^2$ test<br>($p < 0.01$) |
| --- | --- | --- | --- | --- | --- | --- | --- |
| BC<sub>1</sub> Gerek 79 | 340 |  |  |  |  | 174:166 | 0.76<sup>ns</sup> |
| Seed set |  | 0.5 | 2.5 | 0 | 0.03 |  |  |
| Number of sterile<br>spikelets per spike |  | 19.1 | 29.5 | 5.5 | 0.32 |  |  |
| BC<sub>1</sub> 71R1203 | 206 |  |  |  |  | 100:106 | 0.77<sup>ns</sup> |
| Seed set |  | 0.3 | 2.1 | 0 | 0.02 |  |  |
| Number of sterile<br>spikelets per spike |  | 16.8 | 23 | 3.7 | 0.31 |  |  |

ns= not significant

### Construction of genetic maps

Following filtration of 16,762 SNPs used for genotyping of BC_1_ lines, the resulting genetic base maps consisted of 929 and 994 unique SNP loci, spanning 2,160 and 2,328 cM over 21 linkage groups in BC_1_ Gerek 79 and BC_1_ 71R1203, respectively. The average distance (2.4 cM) between two unique loci was similar in both linkage maps (Tables S1 & S2).

Using the categorical fertility phenotypes (completely sterile or fertile), a new restorer locus was mapped as a qualitative (monogenically inherited) trait between SNP markers *IWB72413* (4.3 cM) and *IWB1550* (4.7 cM) in the subtelomeric region of chromosome 6AS in BC_1_ Gerek 79 (Fig. 2a). The newly dissected locus was designated *Rf9* following the Catalogue of Gene Symbols (https://shigen.nig.ac.jp/wheat/komugi/genes/symbolClassListAction.do?geneClassificationId=68) for Restorers for Cytoplasmic Male Sterility in wheat. No restorer locus underlying the binary phenotype could be genetically mapped in BC_1_ 71R1203.

**Fig. 2.**
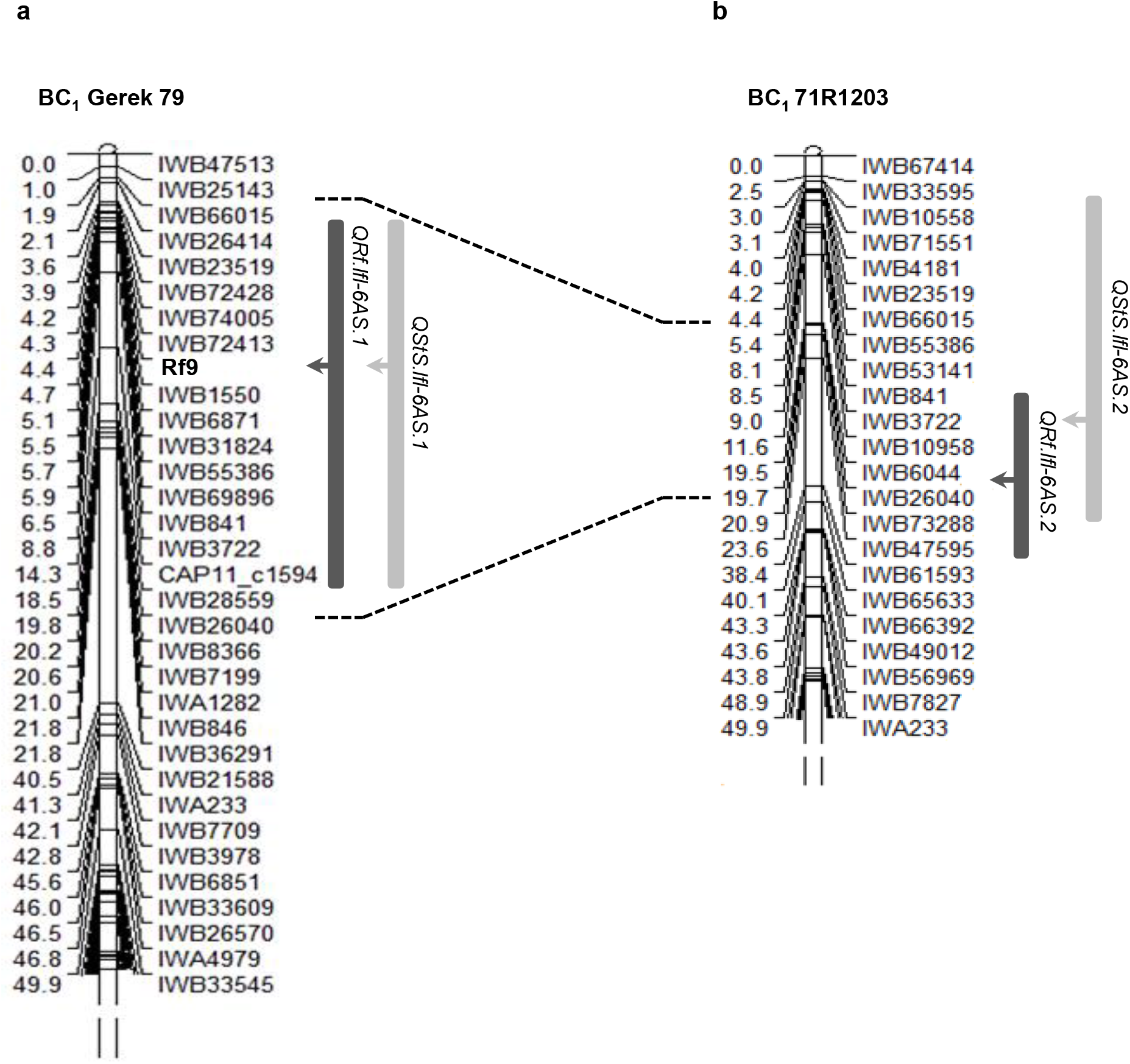
Genetic position of the gene *Rf9* and co-location of QTL detected for seed (*QRf*, dark bars) and number of sterile spikelets per spike *(QStS,* light bars) on chromosome 6AS in (a) BC_1_ Gerek 79 and (b) BC_1_ 71R1203. The peak of each QTL is shown with arrows. Correspondence interval of *IWB66015-IWB26040* common markers in two maps is shown with dot lines. A partial linkage map of the short arm of chromosome 6A is presented.

### Identification of QTL in BC_1_ Gerek 79 and BC_1_ 71R1203

Composite interval mapping in BC_1_ Gerek 79 (Table 3) detected two QTL for seed set on chromosomes 6AS *(QRf.fl-6AS.1)* and 4AL *(QRf.fl-4AL)* that explained 18 % and 14 % of the phenotypic variation, respectively. At both loci, the parental line Sperber contributed with negative additive effects indicating that the Gerek 79 alleles increased seed set values. In this population, three QTL for number of sterile spikelets per spike were identified on chromosomes 6AS *(QStS.lfl-6AS.1),* 6BS *(QStS.lfl-6BS)* and 2AL *(QStS.lfl-2AL).* Of these, *QStS.lfl-6AS.1*, located close to *IWB72428,* showed the highest LOD score (46.3) and explained 53 % of the total phenotypic variation for number of sterile spikelets per spike. The QTL allele that increased the number of sterile spikelets was inherited from the parental line Sperber (Table 3).

**Table 3.** Chromosomal locations, map intervals, flanking markers, LOD scores, additive effects (A.E.) of Sperber and percentage of explained variance by QTL detected for seed set *(QRf)* and number of sterile spikelets per spike (*QStS*) in BC_1_ Gerek 79 and BC_1_ 71R1203.

| Trait/ Population | QTL | Chromosome | Interval (cM) | Flanking markers | LOD | R<sup>2</sup> (%) | A.E. |
| --- | --- | --- | --- | --- | --- | --- | --- |
| BC<sub>1</sub> Gerek 79 |  |  |  |  |  |  |  |
| Seed set | *QRf.lfl-4AL* | 4AL | 48.6-50.7 | *BS00062059-IWB74057* | 3.1 | 14 | -0.46 |
|  | *QRf.lfl-6AS.1* | 6AS | 1.8-14.2 | *IWB66015-CAP11_c1594* | 6.2 | 18 | -0.34 |
| Number of sterile spikelets per spike | *QStS.lfl-2AL* | 2AL | 46.7-47.3 | *IWB12381-IWB53795* | 3.1 | 9 | 2.07 |
|  | *QStS.lfl-6AS.1* | 6AS | 1.8-14.2 | *IWB66015-CAP11_c1594* | 46.3 | 53 | 8.59 |
|  | *QStS.lfl-6BS* | 6BS | 0-34 | *IWA921-IWB12660* | 14 | 21 | 6.35 |
| BC<sub>1</sub> 71R1203 |  |  |  |  |  |  |  |
| Seed set | *QRf.lfl-1AS* | 1AS | 1.6-20.7 | *IWB7436-IWB28549* | 7.2 | 12 | -0.26 |
|  | *QRf.lfl-1BS* | 1BS | 0-10.2 | *IWB69597-IWB12475* | 4.4 | 7 | -0.21 |
|  | *QRf.lfl-5BL* | 5BL | 133.1-147.1 | *IWB33310-IWB26869* | 3.6 | 5 | -0.19 |
|  | *QRf.lfl-6AS.2* | 6AS | 9.0-23.5 | *IWB3722-IWB47595* | 5.9 | 11 | -0.21 |
| Number of sterile spikelets per spike | *QStS.lfl-1AS* | 1AS | 1.6-22.9 | *IWB7436-IWB31602* | 26.3 | 46 | 6.11 |
|  | *QStS.lfl-1DS* | 1DS | 2.5-21.8 | *IWB11524-IWB59019* | 3.6 | 4 | 2.29 |
|  | *QStS.lfl-6AS.2* | 6AS | 2.5-20.8 | *IWB33595-IWB73288* | 5.8 | 12 | 2.73 |

Four QTL for seed set in BC_1_ 71R1203 population (Table 3) were identified on chromosomes 1AS *(QRf.lfl-1AS*), 1BS *(QRf.lfl-1BS),* 5BL *(QRf.lfl-5BL),* and 6AS *(QRf.lfl-6AS.2),* explaining 12, 7, 5 and 11 % of the total phenotypic variation, respectively. A higher seed set was conferred by the 71R1203 allele at all loci. The most significant QTL for number of sterile spikelets per spike with a LOD score of 26.3 was detected on chromosome 1AS *(QStS.fl-1AS)* near to *IWB7436* with a positive additive effect derived from Sperber. This QTL together with two QTL on chromosomes 1DS *(QStS.fl-1DS)* and 6AS *(QStS.fl-6AS.2)* explained 46, 4 and 12 %, respectively, of the total phenotypic variation for the trait (Table 3).

A QTL hotspot for seed set and number of sterile spikelets per spike was deciphered in the nearly 20 cM spanning interval of the common SNP markers *IWB66015-IWB26040* on chromosome 6AS in both mapping populations (Fig. 2a, b). In BC_1_ Gerek 79, the same QTL were identified in the interval between *IWB66015* (1.8 cM) and *CAP11_c1594* (14.2 cM) for controlling seed set *(QRf.lfl-6AS.1)* and number of sterile spikelets per spike *(QStS.lfl-6AS.1)* with the opposite allelic effect of Sperber (−0.34 and 8.59, respectively) (Table 3). Remarkably, the peak of both QTL harboured *Rf9,* located 4.4 cM proximal to the subtelomeric region in BC_1_ Gerek 79 (Fig. 2a). The magnitudes and directions of allelic effects at *Rf9* and SNP loci *IWB72413* (4.3 cM) and *IWB1550* (4.7 cM) showed a highly significant effect for seed set (Fig. 3a) and number of sterile spikelets per spike (Fig. 3b), with the favourable allele derived from Gerek 79. In BC_1_ 71R1203, the QTL for seed set and number of sterile spikelets per spike also overlapped the *IWB66015-IWB26040* interval on chromosome 6AS (Fig. 2b) and revealed negative (−0.21 for *QRf.fl-6AS.2)* and positive (2.73 for *QStS.fl-6AS.2*) allelic effects of Sperber for seed set and number of sterile spikelets per spike, respectively.

**Fig. 3.**
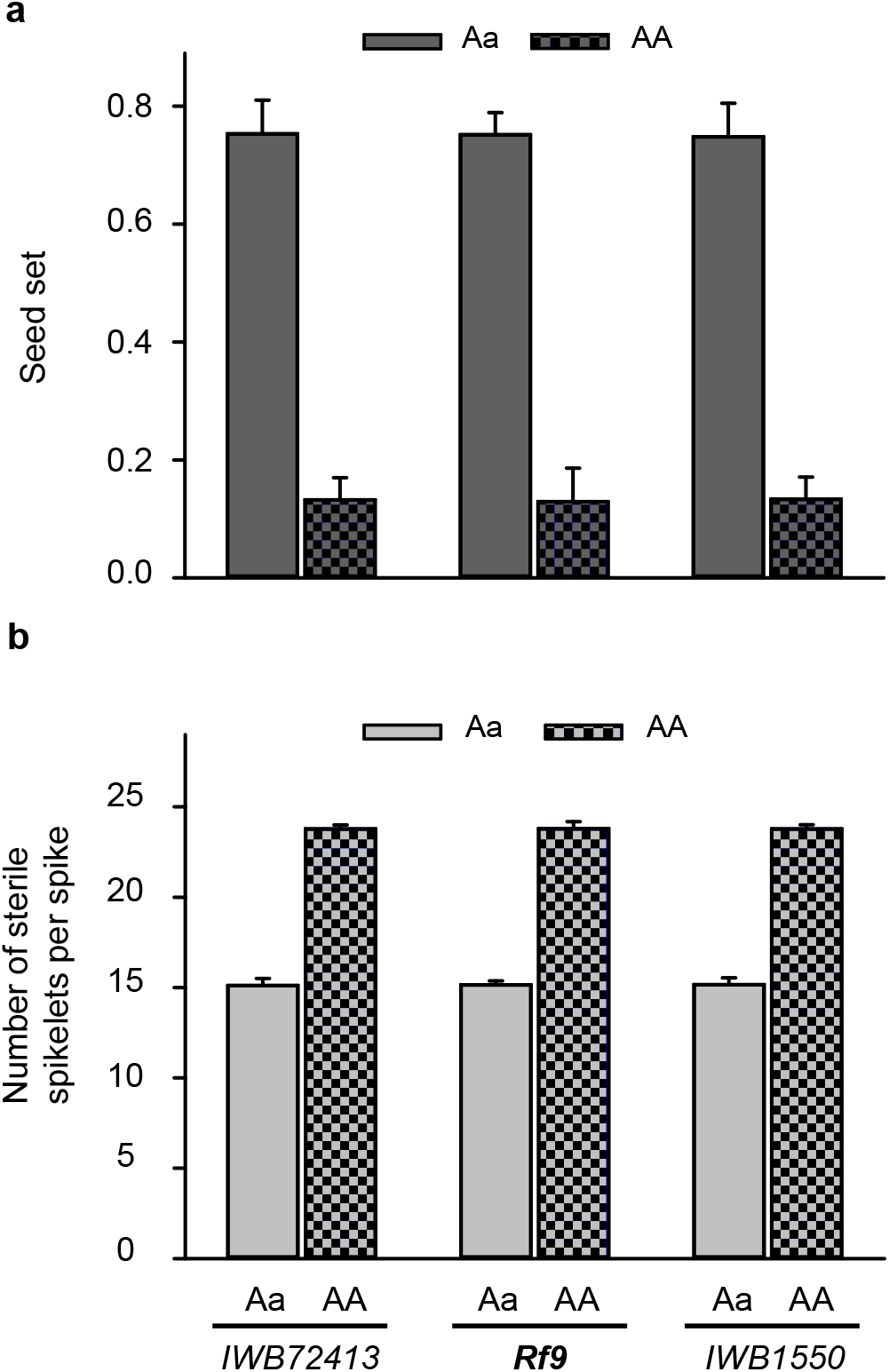
Comparison of allele effects for *Rf9* locus and two flanking SNP loci *(IWB72413* and *IWB1550*) at the peak of *QRf.lfl-6AS.1* and *QStS.lfl-6AS.1* on (a) seed set and (b) number of sterile spikelets per spike in BC_1_ Gerek 79. Dotted bars refer to the alleles of CMS-Sperber (AA) *vs*. restorer line Gerek 79 (Aa). T-tests based on the mean value of each trait showed highly significant differences (*p*<0.001) between the allele effects of CMS-Sperber compared to Gerek 79 parental lines for all loci.

### Candidate genes associated with co-located QTL

To further investigate the chromosomal region associated with co-located QTL controlling the traits on chromosome 6AS, SNP markers *IWB66015* (1.8 cM) and *CAP11_c1594* (14.2 cM) flanking *QRf.lfl-6AS.1* and *QStS.lfl-6AS.1* were used to search for putative candidate genes (Table S3). The search resulted in 93 candidate genes physically located in this interval based on rice annotation translated sequence similarities (RGAP version 7). Among those, 17 genes belonged to the mitochondrial transcription termination factor (mTERF) family. Genes *Traes_6AS_60B99C6F3* encoded for proteins of the tetratricopeptide repeat (TPR) family, whereas *Traes_6AS_97F4D73A5* and *Traes_6AS_A5F88DE9A (Rf1)* encoded pentatricopeptide repeat (PPR) proteins. *In silico* expression analysis of candidate genes (Table S3) showed a wide range of expression for the genes in different organs and at three developmental stages. The gene *Traes_6AS_60B99C6F3* was highly expressed (0.98 FPKM) at Zadoks 39 (flag leaf ligule visible) only in spikes of wheat (Table S3). The expression (Fig. 4) of *Traes_6AS_97F4D73A5* in spikes was higher (1.16 FPKM) at Zadoks 32 (second node visible) whereas *Traes_6AS_A5F88DE9A (Rf1)* was highly expressed (1.99 FPKM) in spikes at Zadoks 65 (full flowering: 50 % of anthers matured). Their expression patterns indicated that they could have biological roles in spike and, more likely spikelet, development in wheat.

**Fig. 4.**
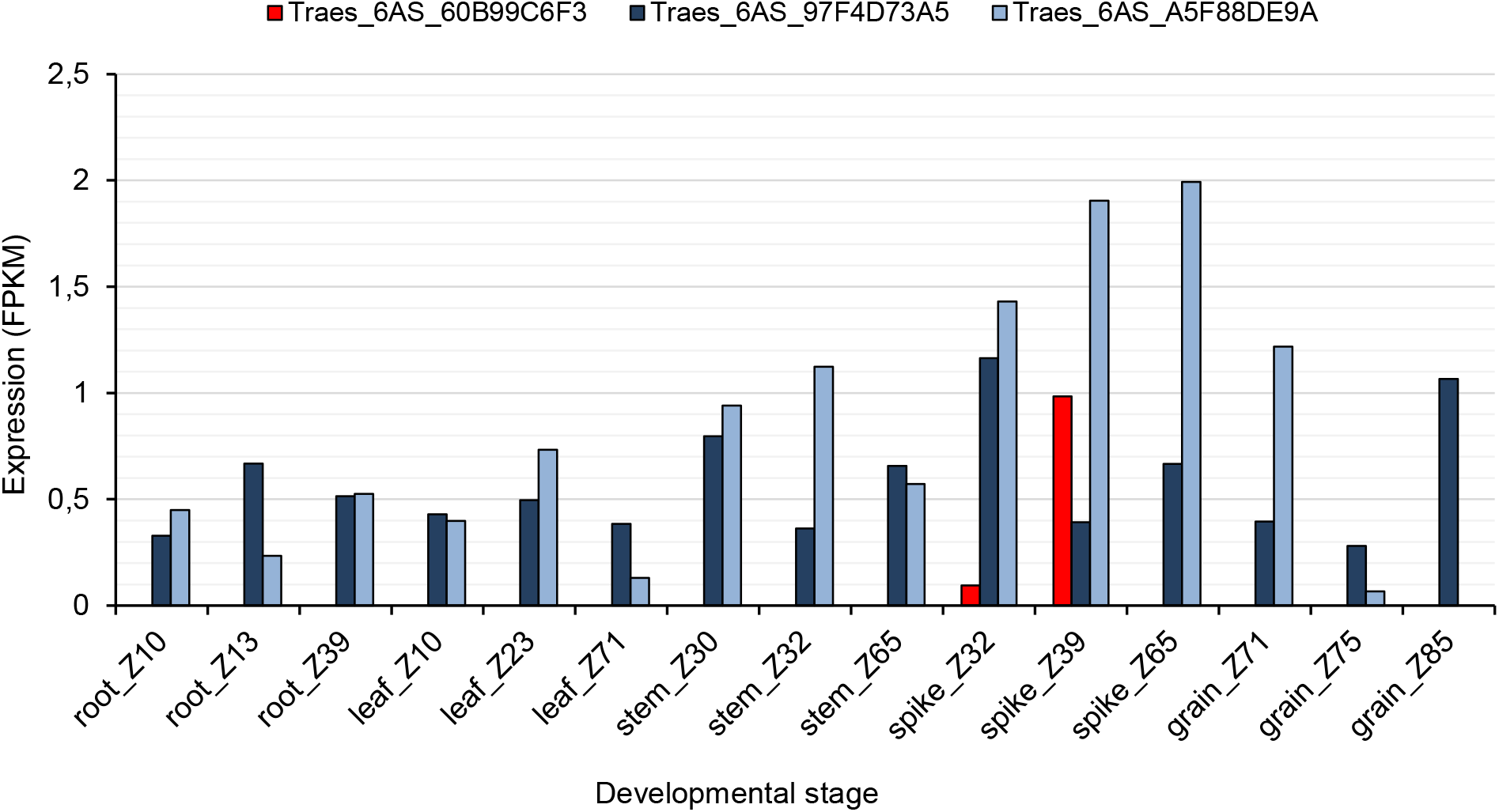
*In silico* expression analysis of the candidate genes *Traes_6AS_60B99C6F3* and *Traes_6AS_97F4D73A5* and *Traes_6AS_A5F88DE9A* (*Rf1*) encoded for tetratricopeptide repeat (TPR) and pentatricopeptide repeat (PPR) containing proteins, respectively in different wheat plant organs at different developmental stages according to Zadoks et al. (1974).

## Discussion

Yield gains associated with heterosis in wheat have been achieved at a slower pace than in other major crops such as maize and rice. The lack of an efficient system for producing hybrid seed is the major bottleneck impairing the competitiveness of hybrid breeding over line-breeding. At the breeding level, it hinders efficient development of large numbers of test crosses required for development of heterotic pools. The CMS hybridisation system based on sterility induced by the cytoplasm of *T. timopheevii* has been proven as a potentially efficient strategy for hybrid seed production. However, further genes and QTL controlling fertility restoration have to be identified and tagged with breeder-friendly molecular markers for a more efficient breeding process. In wheat, this was severely constrained by its large complex genome of about 17 Gb, until the recent release of a reference genome sequence (Appels et al. 2018). QTL for general fertility restoration in common wheat have been identified on chromosomes 1BS, 2AL, 2BS, 4BS and 6AS (Ahmed et al. 2001), 1BS, 5A and 7D (Zhou et al. 2005), 2DS (Dou et al. 2009), 1AS, 1BS and 6BS (Geyer at al. 2018) and 2DS, 4BS and 7AL (Yuan et al. 2020). In the present study, we performed QTL analyses in two populations segregating for restorer lines discovered in Gerek 79 and 71R1203 and identified a novel restorer locus on chromosome 6AS (*Rf9*) together with QTL on chromosomes 1AS, 1BS, 1DS, 2AL, 4AL, 5BL and 6BS.

Gerek 79 was the most widely grown winter wheat cultivar in Turkey in the 1990s. It is a tall wheat derived from a Turkish landrace ‘Yala’ which has resistance to common bunt caused by *Tilletia caries* and *T. foetida* (Bonjean et al. 2001). 71R1203, developed from a cross of winter wheat ‘NB542437’ was one the best restorer lines identified in field trials conducted in Eastern Washington during 1974-1978 (Allan and Rubenthaler 1984). In BC_1_ populations derived from Gerek 79 and 71R1203, we found 12 QTL for seed set and number of sterile spikelets per spike. Four of these QTL *(QRf.fl-6AS.1, QStS.lfl-6AS.1, QRf.lfl-6AS.2, QStS.lfl-6AS.2)* overlapped at the distal end of chromosome 6AS. Ahmed et al. (2001) detected a minor QTL for seed fertility on chromosome 6AS, but located much closer to the centromere using recombinant inbred lines derived from a cross of *T. aestivum* cv. ‘Chinese Spring’ and *T. spelta* var. *duhamelianum*.

It seems the new restorer locus *Rf9* that coincided with the QTL peaks of *QRf.fl-6AS.1* and *QStS.fl-6AS.1* in the subtelomeric region of chromosome 6AS (Fig. 2a) is new with no *Rf* gene previously reported in the region. Putatively annotated high confidence genes located between the flanking markers of the gene *Rf9* showed possible associations with candidate genes coding for TPR, PPR and mTERF proteins with higher expression patterns in the spikes (Table S3). A T-DNA insertion mutant OsAPC6 related to a TPR containing protein in rice caused abnormal mitotic divisions during megagametogenesis and led to reduced plant height and up to 45 % decreased seed set (Awasthi et al. 2012). The involvement of TPR-containing proteins in hybrid sterility was reported by Yu et al. (2016), who showed that down-regulation of ORF3 encoding a TPR domain-containing protein restored spikelet fertility in hybrid rice by sporogametophyte allelic interaction. PPR genes have been cloned and well-characterised as essential components for fertility restoration in rice (Hu et al. 2012) and sorghum (Tang et al. 1996). Members of the mitochondrial transcription termination factor (mTERF) family are also involved in fertility restoration in cereals. Pan et al. (2019) identified the candidate gene *Zea mays small kernel 3 (Zmsmk3),* which contained two mTERF motifs and was required for the intron splicing of mitochondrial nad4 and nad1 and kernel development. Genome-wide association studies in a multiparental mapping population in hybrid barley detected two mTERF proteins linked to the restorer locus *Rfm3* on the short arm of chromosome 6H (Bernhard et al. 2019). Functional association of the wheat *Rf9* locus and these other candidate genes deserves attention in future studies.

In BC_1_ 71R1203, a genome-wide QTL scan revealed two overlapping major QTL, *QRf.lfl-1AS* and *QStS.lfl-1AS,* within the interval *IWB7436* (1.6 cM) – *IWB31602* (22.9 cM) on chromosome 1AS (Fig. 2b). The identified QTL probably corresponded to *Rf1* in line R3, the first reported restorer of *T. timopheevii* CMS (Livers 1964). It was later located on chromosome 1A by monosomic analysis (Yen et al. 1969). Geyer et al. (2018) detected a QTL for seed set in association with fertility restoration *Rf1* locus in a BC_1_ population derived from CMS-Sperber and line R3 restorer.

Composite interval mapping also detected minor QTL *QRf.lfl-4AL, QStS.lfl-2AL and QStS.lfl-6BS* in BC_1_ Gerek 79 and *QRf.lfl-1BS, QRf.lfl-5BL* and *QStS.lfl-1DS*in BC_1_ 71R1203 (Table 3). Except for two QTL on chromosomes 1BS and 6BS, previously identified as minor or modifier loci for major loci *Rf3* and *Rf4* (Ahmed et al. 2001; Zhou et al. 2005; Geyer et al. 2018), we detected new QTL on chromosomes 1DS, 2AL, 4AL and 5BL (Table 3). A KASP assay developed based on the SNP *IWB72107* was used to determine whether QTL *QRf.lfl-1BS* could be associated with *Rf3* located on chromosome 1BS (Geyer et al. 2018). No genetic polymorphism at this SNP locus was found between the parental lines when compared to the marker genotype of Primepi as a reference for the *Rf3* allele (data are not shown), indicating that *Rf3* was not present in BC_1_ Gerek 79 or BC_1_ 71R1203. A comparison of flanking SNP markers and genetic distances also showed that none of these minor QTL was linked to a region associated with a major *Rf* gene detected previously (Ma et al. 1995; Ahmed et al. 2001; Zhou et al. 2005; Stojałowski et al. 2013; Whitford et al. 2013; Lukaszewski 2017).

Both BC_1_ populations exhibited moderate restoration potential leading to incomplete fertility in most of the individuals. Nevertheless, genetic components of restorer parental lines could improve fertility restoration when combined with other restorer loci. Robertson and Curtis (1967) identified modifier genes located on chromosomes 1B, 2A, 3D, 6A and 6B in Chinese Spring wheat that prevented full restoration, even in the presence of *Rf1* and *Rf2.* Geyer et al. (2018) reported a significant effect of modifier loci located on chromosomes 1BS and 6BL on the penetrance of *Rf1,* however; such interactions between major genes and modifier loci were proposed in several studies as essential for full fertility restoration (Ahmed et al. 2001, Zhou et al. 2005; Stojałowski et al. 2013; Würschum et al. 2017). Besides the need to modify the cleistogamous behavior of wheat to a more outcrossing system, the production of commercial hybrid seed will require a stable and efficient hybridization system comprising an easily pollinated female parent and a male parent with ability to completely rerstore fertility under most field conditions. Selection for male characteristics such as plant height, anther size, pollen viability and longevity, general and specific combining ability with the female parent(s) and genetic diversity is important for exploring the best pollinators. However; the emphasis should be on the integration of genetic factors that control fertility restoration comprising QTL and *Rf* genes for developing males through repeated backcrossing. Selection of males segregating for fertility restoration loci in *T. timopheevii* cytoplasm could be performed by test crossing and evaluation of the plants particularly under conditions that might inhibit the expression of fertility restoration such as heat and drought (Bonjean et al. 2001). To reach an acceptable level of hybrid fertility, a combination of two to three genes should be sufficient; therefore restorer lines with effective major QTL and genes such as *Rf1, Rf3,* and *Rf9* could be tested for maximum fertility restoration. Minor QTL for fertility restoration could be used for improving the female lines. Further, the development of functional and predictive molecular markers for tracking fertility restoration loci ultimately increases the efficiency of breeding processes and facilitates production of restorer parents. The results of our study could be used for developing new genetic resources of restorer lines and marker-assisted breeding in hybrid wheat. Also, it provides new insights into the genetic mechanisms controlling fertility restoration in wheat with the cytoplasm of *T. timopheevii* and lays the groundwork for gene characterization, cloning and manipulation in future fundamental studies.

## Acknowledgments

The authors would like to thank Sabine Schmidt, Petra Greim, Mahira Duran and the working group Wheat and Oat Breeding Research of the Bavarian State Research Center for Agriculture for excellent technical assistance. We are thankful to Dr. Bianca Büttner, Dr. Günther Schweizer and Dr. Theresa Albrecht for helpful discussions. We gratefully acknowledge valuable support and cooperation from Dr. Julia Rudloff and Dr. Erhard Ebmeyer, KWS LOCHO, Germany in this project, especially for phenotyping the BC_1_ 71R1203 population. We appreciate Prof. Robert McIntosh for providing critical information on the gene symbols and reviewing the manuscript. The project was supported by funds of the Federal Ministry of Food and Agriculture (BMEL) based on a decision of the Parliament of the Federal Republic of Germany via the Federal Office for Agriculture and Food (BLE) under the innovation support programme.

## Ethical approval

This article does not contain any studies with human participants or animals performed by any of the authors.

## Supplementary Tables

**Table S1** Position of the markers (cM) in 21 linkage groups of wheat in BC_1_ Gerek 79 mapping population.

**Table S2** Position of the markers (cM) in 21 linkage groups of wheat in BC_1_ 71R1203 mapping population.

**Table S3** List of the candidate genes associated with co-located QTL *Q1Rf.lfl-6AS* and *Q1StS.lfl-6AS* including physical position, contig ID, MIPS annotation hit ID and description, rice annotation hit ID and description and expression analysis in wheat organs at different developmental stages according to Zadoks et al. (1974).

